# Single cell RNA sequencing of nc886, a non-coding RNA transcribed by RNA polymerase III, with a primer spike-in strategy

**DOI:** 10.1101/2024.03.20.585884

**Authors:** Gyeong-Jin Shin, Byung-Han Choi, Hye Hyeon Eum, Areum Jo, Nayoung Kim, Huiram Kang, Dongwan Hong, Jiyoung Joan Jang, Hwi-Ho Lee, Yeon-Su Lee, Yong Sun Lee, Hae-Ock Lee

**Author notes:** corresponding authors: YS Lee, (email) H Lee, (email). equal contribution (Gyeong-Jin Shin, Byung-Han Choi).

## Abstract

Single cell RNA sequencing (scRNA-seq) has emerged as a versatile tool in biology, enabling comprehensive genomic-level characterization of individual cells. Currently, most scRNA-seq methods generate barcoded cDNAs by capturing polyA tails of mRNAs, which excludes many non-coding RNAs (ncRNAs), especially those transcribed by RNA polymerase III (Pol III). Although previously thought to be expressed constitutively, Pol III-transcribed ncRNAs are expressed variably in healthy and disease states and play important roles therein, necessitating their profiling at the single cell level. In this study, we have developed a measurement protocol for nc886 as a model case, as an initial step for scRNA-seq for Pol III-transcribed ncRNAs. Specifically, we spiked in an oligo-tagged nc886-specific primer during the polyA tail capture process for the 5’-reading in scRNA-seq. We then produced sequencing libraries for standard 5’ gene expression and oligo-tagged nc886 separately, to accommodate different cDNA sizes and ensure undisturbed transcriptome analysis. We applied this protocol in three cell lines which express high, low, and zero levels of nc886, respectively. Our results show that the identification of oligo tags exhibited limited target specificity, and sequencing reads of nc886 enabled the correction of non-specific priming. These findings suggest that gene-specific primers (GSPs) can be employed to capture RNAs lacking a polyA tail, with subsequent sequence verification ensuring accurate gene expression counting. Moreover, we embarked on an analysis of differentially expressed genes in cell line sub-clusters with differential nc886 expression, demonstrating variations in gene expression phenotypes. Collectively, the primer spike-in strategy allows us for a combined analysis of ncRNAs and gene expression phenotype.

## Introduction

During the past two decades, non-coding RNAs (ncRNAs) and next-generation sequencing (NGS) technologies have been among the greatest advances in biology (1, 2). Numerous studies have documented the diverse biological roles of ncRNAs, with the most prominent ones being the gene-regulatory functions of microRNAs and long ncRNAs (lncRNAs). NGS techniques, which offer unprecedented high-throughput capabilities, have generated enormous amounts of genomic, epigenomic, and transcriptomic data. Additionally, NGS has greatly advanced the field of ncRNAs by enabling the capture of low-copy RNAs, many of which have been identified to be non-coding (3). More recently, NGS has been applied at the single cell level.

Single cell RNA sequencing (scRNA-seq) technologies continue to advance, with the fundamental principle being the creation of barcoded cDNAs that allow for the differentiation of individual cells (4, 5). Particularly, droplet-based approaches such as the Chromium system (10X genomics) provide a combination of simplicity and cost-effectiveness, and they currently constitute the majority of scRNA-seq data (6). A limitation of the system arises from the utilization of oligo-dT sequences during cDNA synthesis, restricting the focus to mRNAs with a polyA tail. Analogous constraints have also been observed in bulk RNA sequencing analysis. Consequently, various strategies are employed to investigate RNA molecules that are typically excluded, such as non-polyAed lncRNAs and small RNAs (7). In bulk RNA sequencing, adaptor ligation or random primers may be applied following the removal of ribosomal RNAs (rRNAs) (8, 9). The addition of polyA or polyU tails also allows for the sequencing of non-polyAed RNAs (10). Several research groups have reported methods for quantifying non-polyAed RNAs at the single cell level (11–13). These methods modified techniques used in total RNA-seq in bulk, including random priming, polyA tailing, and/or rRNA removal. While showing promise, they are developed for C1 microfluidic devices or SMART-seq, which are costly compared with the droplet-based methods.

Despite the increase in research on ncRNAs, a subset of ncRNAs remains underexplored. These are medium-sized ncRNAs that are transcribed by RNA polymerase III (Pol III). They include transfer RNAs (tRNAs), 5S rRNA, and U6 small nuclear RNA. The roles of these RNAs are so fundamental that it is challenging to imagine their dynamic expression. Therefore, Pol III-transcribed ncRNAs (Pol III-ncRNAs) attracted minimal attention during the application and analysis of NGS. However, this view of Pol III-ncRNAs has recently changed. The repertoire of Pol III-ncRNAs are more diverse than previously thought (14). Pol III transcriptomes vary depending on biological situations (15). The best examples are nc886 and tRNA-derived RNA Fragments, which are dynamically expressed and control gene expression (16, 17). Thus, it is essential to obtain Pol III transcriptomes and analyze them in comparison to other -omics data.

As an initial attempt, here we developed a protocol for measuring nc886 in droplet-based scRNA-seq. We chose nc886 for the following reasons (18): Firstly, nc886 is transcribed from a single genomic locus, unlike most other Pol III genes which have identical or highly similar sequences scattered at multiple loci across the genome. Secondly, nc886 has no post-transcriptional modifications, unlike tRNAs. Thirdly, nc886 expression is highly abundant in some cancer cells but is completely silenced in others. These features provide unambiguity in mapping and a set of cell lines for comparison, making nc886 an ideal ncRNA for initially establishing a new sequencing protocol. Furthermore, a single cell expression profile of nc886 will provide valuable information, given its important roles in cancer and immunity

## Materials and methods

### Cell culture, RNA isolation, and qRT-PCR

WPMY-1 and Hep3B cell lines were purchased from the American Type Culture Collection (Manassas, VA). HEK293T was our laboratory stock. We made nc886-expressing Hep3B cells (designated “Hep3B-886” hereafter) by a lentiviral plasmid “pLL3.7.Puro.U6:nc886”. This plasmid was derived from the original lentiviral plasmid, pLL3.7 (Addgene, Watertown, MA), and contains the nc886 gene (a 102 nucleotide (nt)-long DNA fragment) under the U6 promoter (19). Lentivirus production, infection, and selection of puromycin-resistant cells were performed per standard laboratory procedures. From the three cell lines, total RNA was isolated by TRIzol™ Reagent (Invitrogen, Carlsbad, CA) and nc886 was measured by qRT-PCR and Northern hybridization as described previously (20).

### nc886 specific primer design and sequencing library construction

For the generation of a nc886 feature library, a gene-specific primer (GSP) was designed to have 3’ nc886 sequences, flanked by a feature barcode and a sequencing adaptor (Table 1). The sequence is 5’-<u>CGGAGATGTGTATAAGAGACAG</u>NNNNNNNNNN<u>GTATGTCCGCTCGAT</u>NNNNNNNNN<u>AGGGTCAGTAAGCACCCGCG</u>-3’. The first underlined 22 nts represent a Read2N adaptor (10X genomics); the second 15 nts, a feature barcode (TotalSeqTM-C0182, BioLegend); and the third 20 nts, complementary 3’ nc886 sequences. The three nucleotide blocks are separated by 10 or 9 nts spacer sequences. Reverse transcription by this nc886-GSP will generate a 1st strand cDNA consisting of CCC-nc886-spacer-feature barcode-spacer-Read2N sequences. Second strand synthesis will be accomplished by the template switching oligos (TSOs) attached to the gel bead of 5’ scRNA-seq reagent.

### Optimization of Reagent Volumes and Primer Concentration for GEM in nc886 transcript analysis

In the generation of Gel Bead in Emulsion (GEM), we used RT Reagent B, Poly-dT RT Primer, Reducing Agent B and RT Enzyme C, along with nc886-GSP. The reaction contains cell suspension, RT Reagent B (18.8 μl), Poly-dT RT Primer (7.3 μl), Reducing Agent B (1.9 μl), RT Enzyme C (8.3 μl), 10 μM nc886-GSP (0.75 μl, resulting in a final concentration of 0.1 μM), supplemented with nuclease-free water to adjust the total reaction volume to 75 μl.

### 5’ single cell RNA sequencing and read processing

Single cell suspensions of WPMY-1, Hep3B-886, and HEK293T cell lines were mixed in equal numbers and subjected to scRNA-Seq using Chromium Next GEM Single Cell V(D)J Reagent Kits v2 (10× Genomics, Pleasanton, CA). We set the cell recovery rate to 5000 per library and followed the manufacturer’s instructions with a slight modification. During the GEM generation & barcoding step, we added 0.1 µM nc886-GSP to the master mix. In addition to the 5’ gene expression (GEX) library, nc886 feature library was constructed using the 5’ Feature Barcode Kit (10× Genomics). Both libraries were sequenced on an Illumina Hiseq X as 100 bp paired-end. Sequencing reads were mapped to the GRCh38 human reference genome using Cell Ranger toolkit (v5.0.0).

### Processing of oligo-tagged sequences

In raw paired-end reads, preprocessing was performed separately for Read 1 (R1) and Read 2 (R2). For R1, reads containing the nc886 sequence were selected using seqkit (command: seqkit grep -s -p <NC886 sequence>) (21). R2 was filtered for reads containing both the feature barcode sequence and nc886 sequence (command: seqkit grep -s -p <FEATURE barcode sequence> | seqkit grep -s -p <NC886 sequence>). When selecting reads containing the nc886 sequence and allowing mismatches, the number of allowable mismatches was specified using the -m option (e.g., allowing one mismatch: -m 1). After preprocessing R1 and R2 separately, it was possible to encounter cases where the read pairs in R1 and R2 did not match. To address this, a custom python script was used on the preprocessed R1 and R2 to extract only the paired reads. In brief, overlapping read IDs between R1 and R2 were extracted, and the corresponding sequence and quality score information for each ID were extracted to generate new paired reads. These new paired reads were subsequently used in downstream analyses.

### DNA isolation and SNP genotyping array

Genomic DNA was extracted from the three cell lines using PureLinkTM Genomic DNA Mini kit (Invitrogen). Total of 778,783 single nucleotide polymorphisms (SNPs) were genotyped on Infinium Global Screening Array MG v3.0 (Illumina, San Diego, CA) by the local service provider (Macrogen, Seoul, Korea) following the standard Illumina procedures. Normalized signal intensity and genotype were computed using the Illumina/BeadArray Files: Python library. Variant calling format (VCF) genotype file was generated using the GRCh38 reference genome.

### Demultiplexing

To demultiplex three cell line data from pooled scRNA-seq, we followed the freemuxlet (http://github.com/statgen/popscle) workflow (22). Briefly, the popscle tool dsc-pileup was run on the bam file generated by Cell Ranger toolkit and reference vcf file. The reference data was downloaded from Demuxafy (https://demultiplexing-doublet-detectingdocs.readthedocs.io/en/latest/index.html), a supplemental tools that enhances accuracy and subsequent analyses in multiple demultiplexing and doublet detecting methods. Subsequently, the freemuxlet tool, set to its default parameters, was utilized to deconvolve the identities of the sample. Each of three different cell lines (HEK293T, Hep3B-886, WPMY-1) has a distinct VCF file that contains information related to chromosomal positions. The cell lines were distinguished by the similarities between freemuxlet-annotated genotypes and genotypes detected by SNP arrays. During the step, doublets (DBL) and ambiguous (AMB) barcodes are removed (Excluded AMB+DBL : 4,172 cells; HEK293T : 1,687 cells; Hep3B-886 : 2,001 cells; WPMY-1 : 4,738 cells).

### Single cell RNA sequencing analysis using Seurat

From the Cell Ranger outputs, raw gene-cell-barcode matrix was processed using Seurat v4.2.2 R package (23). Low-quality cells were filtered with the criteria nCount>2000 and percent.mito <15. Potential multiplets were predicted by Scrublet and removed (24). After the QC filtering process, the unique molecular identifier (UMI) count matrix was log-normalized and scaled by z-transform. Utilizing the PC ElbowPlot functions of Seurat, PC 7 was selected as a distinct subset of principal components. Subsequently, cell clustering and Uniform Manifold Approximation and Projection (UMAP) visualization were conducted using the ‘FindClusters’ and ‘RunUMAP’ functions. The resolution was set to 0.3 or 0.6, segregating three or six clusters respectively.

### Pathway enrichment analysis and data visualization

Subcluster analysis for WPMY-1 cells was conducted using the ‘enrichGO’ function of the ‘clusterProfiler’ R package (version 4.6.2), focusing on the top differentially expressed genes (DEGs). Genes were filtered based on the adjusted p-value and q-value (< 0.05). The ‘org.Hs.eg.db’ annotation package (version 3.15.0) was utilized for organism-specific categorization. Data filtering was applied as GeneRatios greater than 0.10, and the results were organized in ascending order of adjusted p-values, specifically targeting the ‘biological process’ category in the Gene Ontology.

## Results

### Generation of nc886 feature library using an oligo-tagged gene-specific primer

To test the feasibility of using a GSP during droplet-based scRNA-seq procedures (10x genomics chromium system), we selected nc886 as the model gene and three cell lines with different levels of nc886 expression (Fig 1A). To capture nc886 transcripts which have no polyA tail, we used a GSP with additional feature barcode and adaptor sequences (Fig 1B). The modifications implemented in the GEM generation and the Barcoding reaction mix are detailed in the Methods section. Addition of the GSP allows extension of the nc886 transcript, yielding cDNA containing the nc886 sequence flanked by adaptors to enable library construction. According to our design, the resulting nc886 feature barcode library is expected to contain Read 1 sequences, 10x cell barcode, UMI, and TSO at the 5’ end as well as 15 nts-feature barcode and ‘Read 2’ sequences. In parallel, a 5’ scRNA-seq library for gene expression analysis was produced as a separate sequencing material.

**Fig 1.**
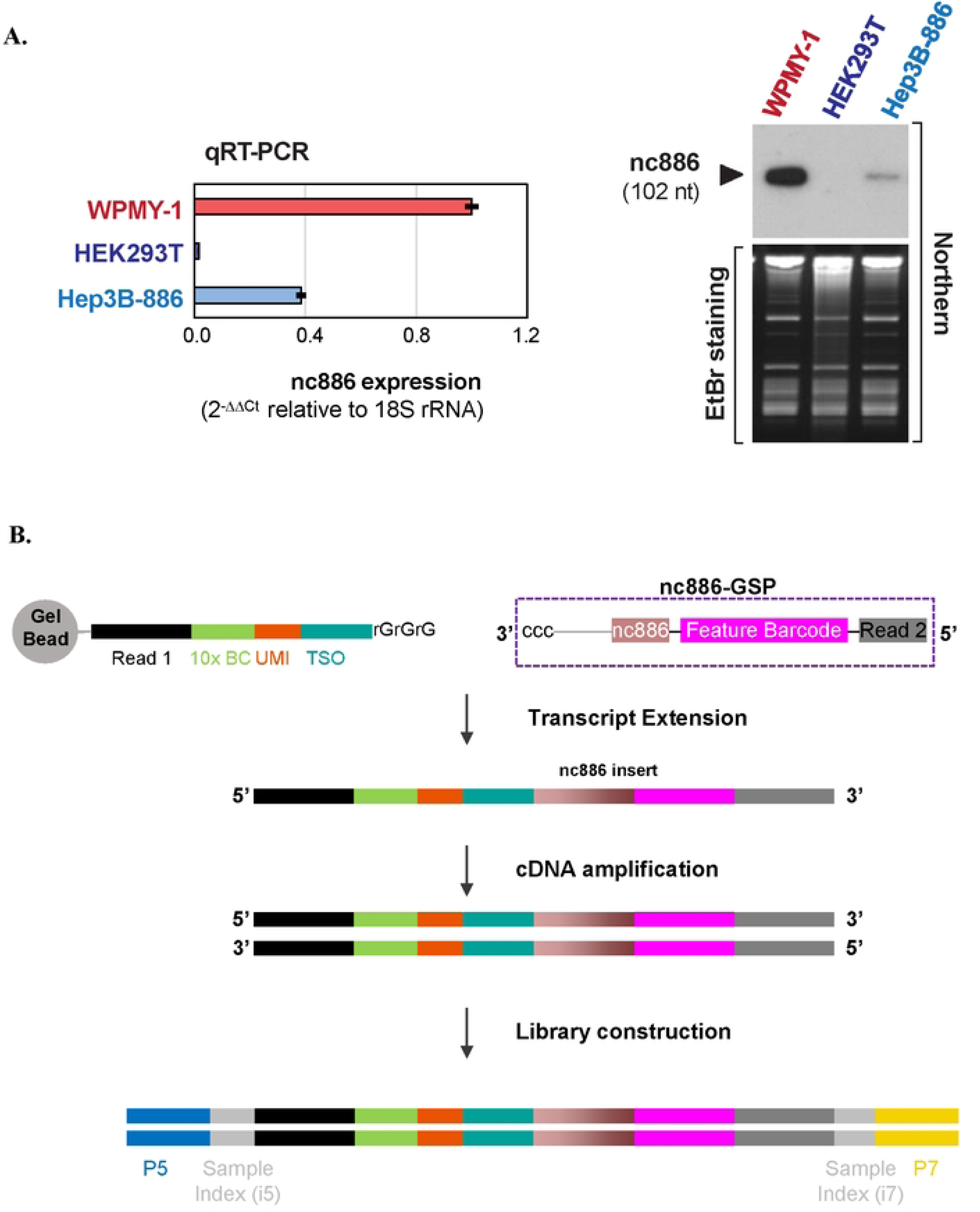
Modification of 5’ scRNA-seq for nc886 detection. **(A)** nc886 gene expression levels in three cell lines, measured by qRT-PCR (left panel) and by Northern hybridization (right panel). In qRT-PCR, each bar represents an average of triplicate samples, with the standard deviation indicated. **(B) A** cartoon depicting the procedure for library preparation in which nc886 gene-specific primer (nc886-GSP) was spiked-in. Diagrams are drawn to show nc886-GSP and products (whose actual sequences are listed in Table 1) in each step. The final library contains sample index and sequencing adaptors PS and P7.

### Determining cell line identities using SNP and gene expression profiles

In this study, we pooled three cell lines with differential nc886 expression to generate multiplex data. WPMY-1 is a myofibroblast cell line derived from a prostate cancer patient (25). Hep3B-886 was derived from a liver cancer cell line, Hep3B, with epithelial morphology and hepatitis B virus integration (26). The HEK293T cell line originated from the human embryonic kidney (27, 28). The first step in our data analysis was to systematically assign pooled scRNA-seq data into respective cell lines using SNP patterns. This segregation was a prerequisite for the investigation of phenotypic alterations in gene expression, especially regarding nc886 expression levels. We utilized Freemuxlet, a tool recommended by the 10x Genomics Analysis Guide (https://www.10xgenomics.com/resources/analysis-guides/bioinformatics-tools-for-sample-demultiplexing), to categorize cells into three distinct clusters. Overlap between SNPs detected in genomic DNA and scRNA-seq data provided cell line identity for each SNP cluster (Fig 2A). Cells corresponding to doublets and ambiguous categories in SNP expression were excluded from subsequent analyses.

**Fig 2.**
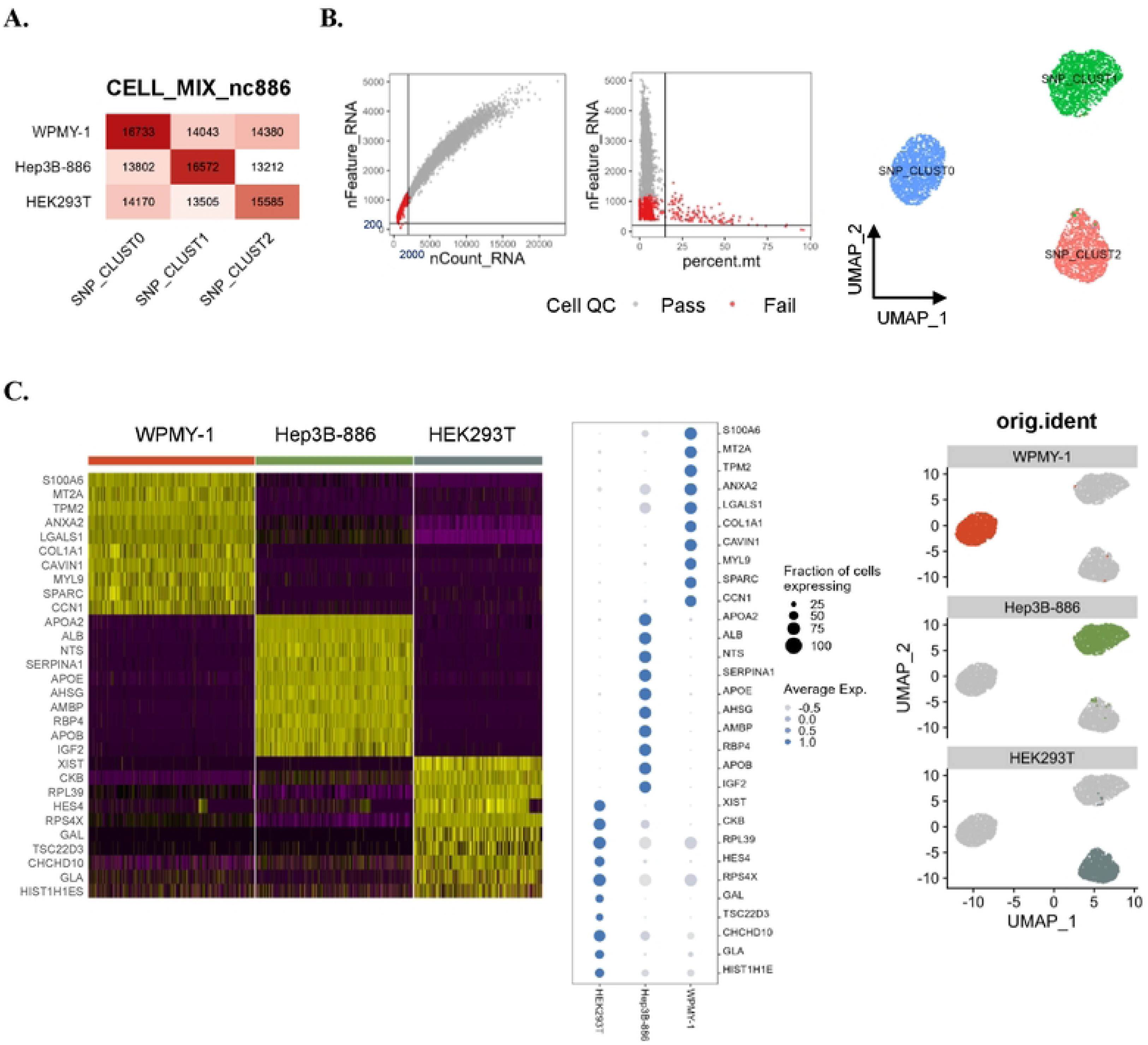
Gene expression profiling of each cell line. **(A)** Cells were classified into three distinct clusters using ’freen1uxlet’. Number of overlapping SNPs between the clusters and SNP arrays for each cell line are indicated. Based on the SNP expression, doublets and ambiguous cells are excluded. **(B)** After freemuxlet runs and doublet/ambiguous cell removal, additional QC filtration was applied: UMI counts(nCount_RNA>2,000), number of genes expressed (nFeature_RNA>200), and propo11ions of mitochondrial gene expression (percent.mt<IS) (left). The UMAP shows SNP clusters within 3 Seurat clusters: SNP_CLUSTO =“WPMY-1”, SNP_CLUSTl =“Hep3B-886”, SNP_CLUST2 = “HEK293T” (right). **(C)** DEG analysis showing gene expression characteristics for each cell line, shown as Heatmap (left), DotPlot (middle). UMAP cluster designation to each cell line (right).

In the second step, we performed clustering analysis based on the 5’ gene expression after further quality control (QC) filtration to select cells with a minimum UMI count of 2,000, a minimum gene count of 200, and a maximum mitochondrial gene proportion of 15 % (Fig 2B, left). Adhering to these criteria, we obtained three clusters assigned as WPMY-1 cells (1,886 cells), Hep3B-886 (1,782 cells), and HEK293T (1,457 cells) (CLUST0, 1, and 2 respectively in the right panel of Fig 2B). Comparison of DEGs in each cluster revealed gene expression characteristics of the three cell lines (Fig 2C). Cluster 0 showed prominent expression of mesenchymal genes such as COL1A1, SPARC and CCN1, which characterize WPMY-1 cells of myofibroblast origin (29). In the Hep3B-886 cluster, liver-specific genes such as ALB, RBP4 and AHSG were highly expressed (30, 31). In the cluster of nc886-silenced HEK293T cells, high expression levels of XIST, TSC22D3 and RPS4X were noted. These expression patterns were consistent with those observed in the original parental cell line (32). These gene expression characteristics confirmed the successful implementation of multiplexing and demultiplexing strategies in scRNA-seq analysis.

### Assessing nc886 gene expression using feature barcoding and sequence alignment strategies

Next, we estimated nc886 expression levels using feature barcode expression data (Fig 3A). However, this method resulted in unexpectedly high nc886 expression in the silenced HEK293T cell cluster (Fig 3B, left). This inconsistency might have been caused by limited primer specificity in our initial feature barcoding approach. To apply higher stringency during mapping nc886 sequences, we analyzed Read 2 (see Fig 1B) from the feature barcode library and specifically extracted nc886 reads (Fig 3A). Using nc886-specific sequences from the untrimmed feature barcode data dramatically reduced the total number of reads (Fig 3A, right), indicating that non-specific priming indeed occurred. The problem of reduction of read numbers was solved by the allowance of a single nucleotide mismatch. This rectified procedure, extraction of nc886 sequence from Read 2 with up to 1 nt mismatch, yielded a result aligned well with the known nc886 expression levels: they were markedly higher in WPMY-1, lower in Hep3B-886, and absent in HEK293T (Fig 3B, right).

**Fig 3.**
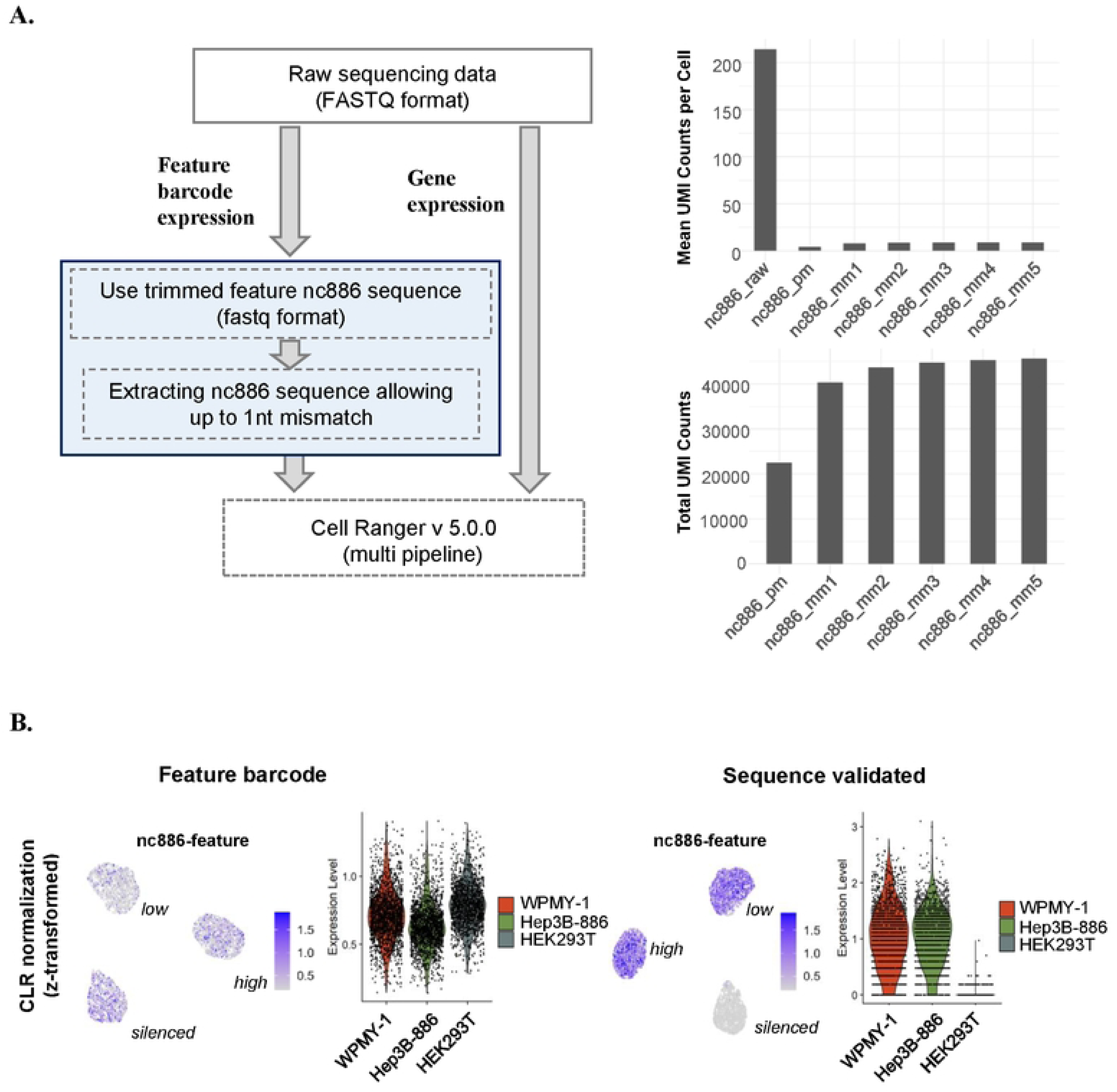
Assessment of nc886 expression level across the cell lines, employing feature barcoding and a sequence alignment strategy. **(A)** Overview of the estimation of nc886 gene expression in comparison to transcriptome analysis (left). To assess nc886 gene expression, the feature barcode or nc886 sequence aligned reads were counted. nc886 gene expression and transcriptome data were processed by the CellRanger rnulti pipeline. Bar plots quantify UMI counts for the feature barcode or nc886 sequence alignn1ents, with n1ismatch counts ranging from one to five nts:“nc886_pm” and “nc886_mm1-5” denote perfect match to nc886 and the number of mismatches (mm) respectively (right). **(B)** The expression level of nc886 across cell lines in z-transform values of the feature barcode (left) or nc886-aligned counts (right) using ’FeaturePlot’ and ’ViolinPlot’.

### Clustering analysis demonstrating diversity in gene expression patterns and nc886 levels

To determine whether each cell line shows heterogeneity in gene expression and nc886 levels, we re-performed clustering analysis with a higher resolution setting. Clustering in the UMAP space revealed the presence of two distinct clusters for each cell line (Fig 4A, upper UMAPs and a heat map). Subsequently, we checked whether the cluster separation reflects differential cell cycle phases (Fig 4A, lower UMAP and bar graph). Hep3B-886 (Hep 1 and Hep 2) and HEK293T cell (HEK 1 and HEK 2) clusters showed different cell cycle distribution between clusters. In contrast, WPMY-1 clusters (W1 and W2) manifested similar cell cycle phases. Thereafter, we performed DEG analysis using the Wilcoxon Rank Sum test (Fig 4A, right), with a particular focus on the WPMY-1 cell line. Comparison of nc886 expression levels between W1 and W2 clusters showed enrichment of nc886 high cells in the W2 cluster (Fig 4B). In the W2 cluster, DEGs include POSTN, MFAP4, DCN, and LUM (Fig 4C), which are closely involved in the extracellular matrix organization as well as in cancer invasiveness (33, 34). By comparison, the W1 cluster DEGs contained CAV1, MT2A, KCNMA1, and CCND1 curated in the response to the metal ion pathway as well as FABP5, SPHK1, and CCN in the regulation of lipid metabolic process (35–38). Consistently, Gene Set Enrichment Analysis (GSEA) annotated protein folding and stability for W1 DEGs and extracellular matrix organization and stimulus for W2 DEGs (Fig 4D). Overall, this combined analysis of gene expression phenotype and nc886 levels suggested that nc886 expression are variable among cells and that this variability contributed to the functional heterogeneity among cells.

**Fig 4.**
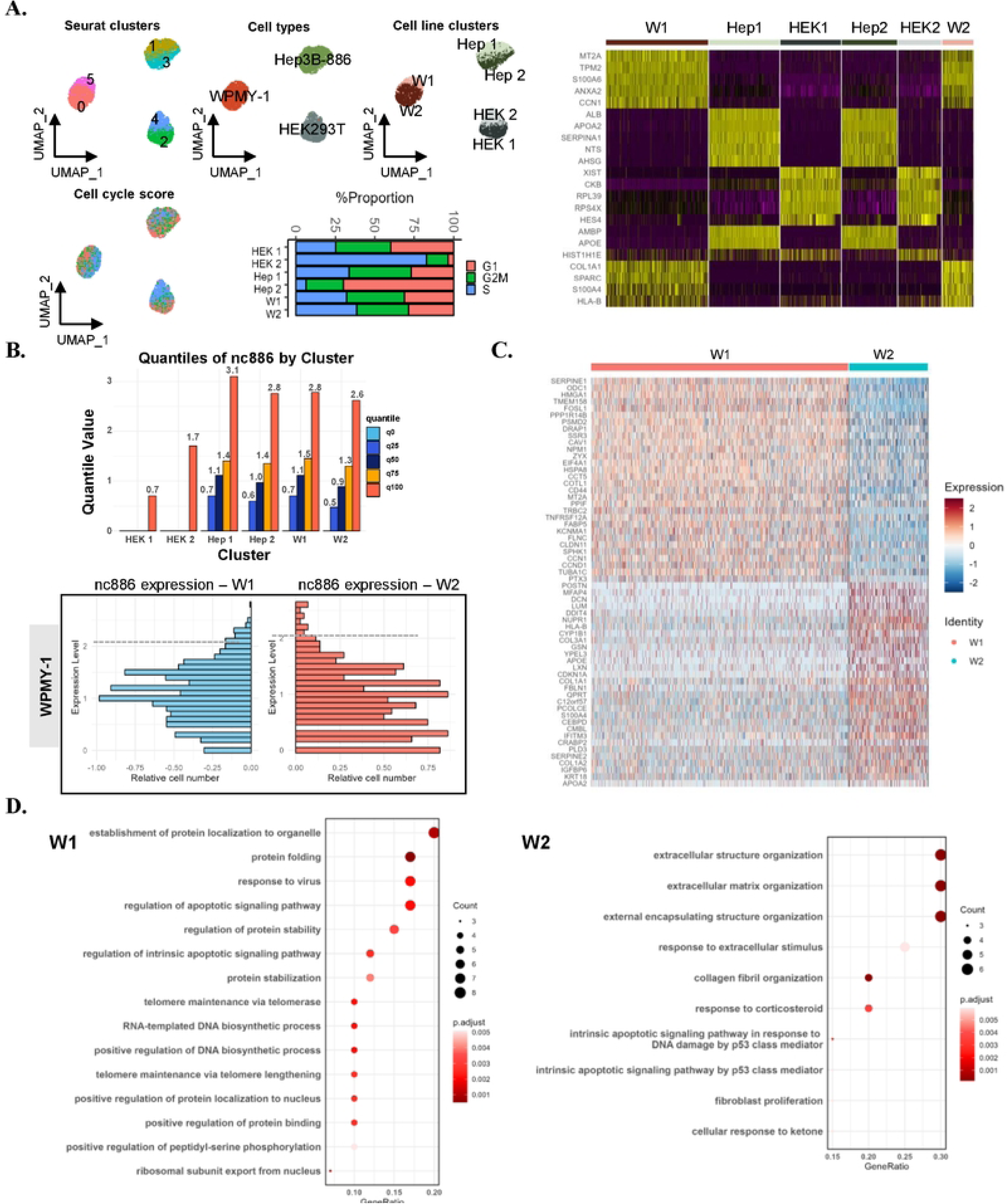
Phenotypic features and nc88^-^6 ^-^ex^-^pression levels in sub-clusters in each of the cell lines. **(A)** UMAP visualization of two distinct clusters per cell line and cell cycle scores. Bar plot; Proportion of cell cycle score distribution in each of the 6 clusters (left). (Cluster 0, **WI;** Cluster I, Hep I; Cluster 2, HEK I; Cluster 3, Hep2; Cluster 4, HEK2; Cluster 5, W2). A heatmap visualization of cluster-specific DEGs obtained from the Wilcox test (right) **(B)** Distribution of nc886 expression analyzed using Quantile and Histogram in the cell line clusters. **(C)** A heatmap showing gene expression of two clusters, WI and W2, in the WPMY-1 cell line. **(D)** GSEA performed on the WPMY-1 clusters, ordered by Gene ratio and adjusted p-value. (cutoff GeneRatio >0.15, p-value <0.005, q-value <0.05).

## Discussion

### Improving nc886 detection in Feature Barcoding: Overcoming inefficiency and non-specific binding

In our study, we used a Feature Barcoding technology to capture nc886, an ncRNA lacking a polyA tail. However, our procedure yielded non-specific sequences in addition to nc886. Although we were able to filter out nc886 sequences and perform subsequent analysis, we should conclude that scRNA-seq of nc886 had limited specificity and was not as effective. A significant proportion of the non-specifically captured sequences were rRNAs.

The main reason for this limitation may be the intrinsic fact that nc886 is a short RNA transcribed by Pol III. This feature made the design of a GSP very challenging. Most Pol III genes have 4-6 consecutive thymidylates at the 3’ end. As a type 2 Pol III gene, nc886 has two intragenic promoter elements, box A and B, each about 15 nts long. Therefore, >30% of the nc886 sequence is potentially homologous to several hundred type 2 Pol III genes. A GSP must lie outside these common sequence motifs and should be located at the 3’ side. The design of a satisfactory primer was very limited and it was almost impractical to design several primers and select the best one. The nc886-GSP used here, which was inevitably designed, may not have been a suitable one. Although it was predicted to be specific in silico, the primer may have been inefficient in recognizing nc886, possibly because of its secondary structures (39). The excessive use of additional sequences in the nc886-GSP may have contributed to compromising specificity in favor of binding to highly abundant rRNAs.

To overcome the limitation found in this study, several improvement strategies can be considered. First, the use of rRNA depletion methods may improve the results since a significant portion of the non-specific sequences were rRNA sequences. Second, cDNA synthesis at high temperature with a thermophilic reverse transcriptase might be a solution for the secondary structure problem. In addition, utilizing a hybridization approach rather than primer extension could potentially yield better results in this non-specific issue. Fixed RNA Seq method developed by the 10X genomics employs the hybridization technique for the gene expression analysis, which may be adopted for the detection of nc886. However, the limited choices in primer design still remain a major obstacle when we recall that our goal in this nc886 study was to lay the groundwork for other Pol III genes and ultimately to obtain single cell Pol III transcriptomes.

### Impact of nc886 on the phenotypic characteristics of cell lines through cluster examination

In this study, we employed SNP data for cluster-based classification of cell lines, aiming to investigate the impact of nc886 expression levels on gene expression patterns.

Within a cell line, we expected nc886 expression levels to be highly variable among individual cells because nc886 has a short half-life (1∼2 hours) and its expression is affected by growth conditions. We observed that nc886 levels became low when we cultured cells in low serum or at high density (YSL, unpublished data). Thus, local variation in cell density would result in different nutrient status of individual cells. Indeed, we observed variable expression levels of nc886 in our scRNA-seq data on WPMY-1 (Fig 4B). We speculate that this difference is not due to a clonal character of each cell, but reflects a temporally transient variation of each cell. This transient variation appeared to have a marginal effect on gene expression, based on our data that two DEG clusters did not show a large difference in nc886 expression levels.

Moving to actual samples, such as those from colon cancer, may reveal further differences in gene expression phenotypes and functionality related to nc886 expression. We aim to improve the methods we have conducted to apply scRNA-seq analysis to samples with higher cellular and functional heterogeneity.

## Acknowledgements

This research was supported by the National Research Foundation of Korea (NRF) funded by the Ministry of Science, ICT & Future Planning (RS-2023-00220840 to HL) and by grants from the National Cancer Center, Korea (NCC-2210320 and NCC-2311362 to YSL and NCC-2210360 to Y-SL). We also acknowledge the Basic Medical Science Facilitation Program, through the Catholic Medical Center of the Catholic University of Korea funded by the Catholic Education Foundation and the KREONET/GLORIAD service provided by KISTI (Korea Institute of Science and Technology Information).

## Notes

### Competing Interest Statement

The authors have declared no competing interest.

## References

1. Cech TR, Steitz JA. The noncoding RNA revolution-trashing old rules to forge new ones. Cell. 2014;157(1):77–94.

2. van Dijk EL, Jaszczyszyn Y, Naquin D, Thermes C. The Third Revolution in Sequencing Technology. Trends Genet. 2018;34(9):666–81.

3. Djebali S, Davis CA, Merkel A, Dobin A, Lassmann T, Mortazavi A, et al. Landscape of transcription in human cells. Nature. 2012;489(7414):101–8.

4. Haque A, Engel J, Teichmann SA, Lonnberg T. A practical guide to single-cell RNA-sequencing for biomedical research and clinical applications. Genome Med. 2017;9(1):75.

5. Baysoy A, Bai Z, Satija R, Fan R. The technological landscape and applications of single-cell multi-omics. Nat Rev Mol Cell Biol. 2023;24(10):695–713.

6. Liao Y, Raghu D, Pal B, Mielke LA, Shi W. cellCounts: an R function for quantifying 10x Chromium single-cell RNA sequencing data. Bioinformatics. 2023;39(7).

7. Kukurba KR, Montgomery SB. RNA Sequencing and Analysis. Cold Spring Harb Protoc. 2015;2015(11):951–69.

8. Cui P, Lin Q, Ding F, Xin C, Gong W, Zhang L, et al. A comparison between ribo-minus RNA-sequencing and polyA-selected RNA-sequencing. Genomics. 2010;96(5):259–65.

9. Huang R, Jaritz M, Guenzl P, Vlatkovic I, Sommer A, Tamir IM, et al. An RNA-Seq strategy to detect the complete coding and non-coding transcriptome including full-length imprinted macro ncRNAs. PLoS One. 2011;6(11):e27288.

10. Li X, Yu K, Li F, Lu W, Wang Y, Zhang W, et al. Novel Method of Full-Length RNA-seq That Expands the Identification of Non-Polyadenylated RNAs Using Nanopore Sequencing. Anal Chem. 2022;94(36):12342–51.

11. Sheng K, Cao W, Niu Y, Deng Q, Zong C. Effective detection of variation in single-cell transcriptomes using MATQ-seq. Nat Methods. 2017;14(3):267–70.

12. Isakova A, Neff N, Quake SR. Single-cell quantification of a broad RNA spectrum reveals unique noncoding patterns associated with cell types and states. Proc Natl Acad Sci U S A. 2021;118(51).

13. Kouno T, Carninci P, Shin JW. Complete Transcriptome Analysis by 5’-End Single-Cell RNA-Seq with Random Priming. Methods Mol Biol. 2022;2490:141–56.

14. Dieci G, Fiorino G, Castelnuovo M, Teichmann M, Pagano A. The expanding RNA polymerase III transcriptome. Trends Genet. 2007;23(12):614–22.

15. Yeganeh M, Hernandez N. RNA polymerase III transcription as a disease factor. Genes Dev. 2020;34(13-14):865–82.

16. Lee YS, Lee YS. nc886, an RNA Polymerase III-Transcribed Noncoding RNA Whose Expression Is Dynamic and Regulated by Intriguing Mechanisms. Int J Mol Sci. 2023;24(10).

17. Wilson B, Dutta A. Function and Therapeutic Implications of tRNA Derived Small RNAs. Front Mol Biosci. 2022;9:888424.

18. Park JL, Lee YS, Song MJ, Hong SH, Ahn JH, Seo EH, et al. Epigenetic regulation of RNA polymerase III transcription in early breast tumorigenesis. Oncogene. 2017;36(49):6793–804.

19. Lee K, Kunkeaw N, Jeon SH, Lee I, Johnson BH, Kang GY, et al. Precursor miR-886, a novel noncoding RNA repressed in cancer, associates with PKR and modulates its activity. RNA. 2011;17(6):1076–89.

20. Kunkeaw N, Lee YS, Im WR, Jang JJ, Song MJ, Yang B, et al. Mechanism mediated by a noncoding RNA, nc886, in the cytotoxicity of a DNA-reactive compound. Proc Natl Acad Sci U S A. 2019;116(17):8289–94.

21. Shen W, Le S, Li Y, Hu F. SeqKit: A Cross-Platform and Ultrafast Toolkit for FASTA/Q File Manipulation. PLoS One. 2016;11(10):e0163962.

22. Kang HM, Subramaniam M, Targ S, Nguyen M, Maliskova L, McCarthy E, et al. Multiplexed droplet single-cell RNA-sequencing using natural genetic variation. Nat Biotechnol. 2018;36(1):89–94.

23. Stuart T, Butler A, Hoffman P, Hafemeister C, Papalexi E, Mauck WM, 3rd, et al. Comprehensive Integration of Single-Cell Data. Cell. 2019;177(7):1888–902 e21.

24. Wolock SL, Lopez R, Klein AM. Scrublet: Computational Identification of Cell Doublets in Single-Cell Transcriptomic Data. Cell Syst. 2019;8(4):281–91 e9.

25. Webber MM, Trakul N, Thraves PS, Bello-DeOcampo D, Chu WW, Storto PD, et al. A human prostatic stromal myofibroblast cell line WPMY-1: a model for stromal-epithelial interactions in prostatic neoplasia. Carcinogenesis. 1999;20(7):1185–92.

26. Aden DP, Fogel A, Plotkin S, Damjanov I, Knowles BB. Controlled synthesis of HBsAg in a differentiated human liver carcinoma-derived cell line. Nature. 1979;282(5739):615–6.

27. Subedi GP, Johnson RW, Moniz HA, Moremen KW, Barb AW. High Yield Expression of Recombinant Human Proteins with the Transient Transfection of HEK293 Cells in Suspension. J Vis Exp. 2015(106):e53568.

28. Tan E, Chin CSH, Lim ZFS, Ng SK. HEK293 Cell Line as a Platform to Produce Recombinant Proteins and Viral Vectors. Front Bioeng Biotechnol. 2021;9:796991.

29. Yu L, Wang CY, Shi J, Miao L, Du X, Mayer D, et al. Estrogens promote invasion of prostate cancer cells in a paracrine manner through up-regulation of matrix metalloproteinase 2 in prostatic stromal cells. Endocrinology. 2011;152(3):773–81.

30. Segal JM, Kent D, Wesche DJ, Ng SS, Serra M, Oules B, et al. Single cell analysis of human foetal liver captures the transcriptional profile of hepatobiliary hybrid progenitors. Nat Commun. 2019;10(1):3350.

31. Sun N, Lee YT, Zhang RY, Kao R, Teng PC, Yang Y, et al. Purification of HCC-specific extracellular vesicles on nanosubstrates for early HCC detection by digital scoring. Nat Commun. 2020;11(1):4489.

32. Wu C, Macleod I, Su AI. BioGPS and MyGene.info: organizing online, gene-centric information. Nucleic Acids Res. 2013;41(Database issue):D561–5.

33. Buechler MB, Pradhan RN, Krishnamurty AT, Cox C, Calviello AK, Wang AW, et al. Cross-tissue organization of the fibroblast lineage. Nature. 2021;593(7860):575–9.

34. Zhu K, Cai L, Cui C, de Los Toyos JR, Anastassiou D. Single-cell analysis reveals the pan-cancer invasiveness-associated transition of adipose-derived stromal cells into COL11A1-expressing cancer-associated fibroblasts. PLoS Comput Biol. 2021;17(7):e1009228.

35. Wang L, Yin YL, Liu XZ, Shen P, Zheng YG, Lan XR, et al. Current understanding of metal ions in the pathogenesis of Alzheimer’s disease. Transl Neurodegener. 2020;9:10.

36. Ling XB, Wei HW, Wang J, Kong YQ, Wu YY, Guo JL, et al. Mammalian Metallothionein-2A and Oxidative Stress. Int J Mol Sci. 2016;17(9).

37. Seo J, Jeong DW, Park JW, Lee KW, Fukuda J, Chun YS. Fatty-acid-induced FABP5/HIF-1 reprograms lipid metabolism and enhances the proliferation of liver cancer cells. Commun Biol. 2020;3(1):638.

38. Zhao L, Wang Z, Xu Y, Zhang P, Qiu J, Nie D, et al. Sphingosine kinase 1 regulates lipid metabolism to promote progression of kidney renal clear cell carcinoma. Pathol Res Pract. 2023;248:154641.

39. Calderon BM, Conn GL. Human noncoding RNA 886 (nc886) adopts two structurally distinct conformers that are functionally opposing regulators of PKR. RNA. 2017;23(4):557–66.

